# Diffusion-based size determination of solute particles: a method adapted for postsynaptic proteins

**DOI:** 10.1101/2025.02.05.636588

**Authors:** András László Szabó, Eszter Nagy-Kanta, Soma Varga, Edit Andrea Jáger, Csaba István Pongor, Mária Laki, András József Laki, Zoltán Gáspári

## Abstract

The postsynaptic density (PSD) is a complex, multi-layered protein network largely situated on the internal surface of the postsynaptic membrane. It is the first processing unit for incoming synaptic transmissions, and changes in its internal structure are associated with synaptic strength and plasticity. These structural changes are largely governed by multivalent interactions between its components. The *in vitro* characterization of such complexes requires unbiased methods that can be used to estimate the size of the emerging assemblies for systems with multiple possible stoichiometries. Here we present an experimental method for detecting specific PSD proteins as well as their complexes based on their diffusion in a microfluidic environment. The method requires a fluorescent labelling technique that does not disrupt the function of labelled proteins, a microfluidic device that can maintain laminar flow for protein solutions, a microscope that can record the fluorescent signal emitted by these solutions, and an analytic software package that can process the collected experimental data and convert them into approximate particle sizes. We demonstrate the applicability of our method on protein constructs of various postsynaptic proteins, including the multivalent assembly between GKAP and LC8.

## Introduction

### Size determination of dynamic protein complexes

Investigation of protein-protein interactions often relies on the characterization of simplified systems. With the exception of relatively small globular proteins, in many cases only specific segments of one or both partners are used. However, a large portion of human proteins contain long disordered segments, often with multiple binding sites, and the exact size, shape, and composition of physiologically occurring complexes is determined by the interplay between the individual binding events to these. [1] In multivalent scenarios, binding between the same partners might occur at different sites, and there might be a dynamic interchange between the individual interactions, in extreme cases leading to the emergence of so-called fuzzy complexes. [2] The stoichiometry of the emergent multivalent complexes might also be variable, depending on the presence of additional structural constraints. Experimental characterization of such dynamic interactions that might give rise to heterogeneous molecular populations requires methods that provide an estimate of the complex size(s) without *a priori* hypotheses, work well in aqueous solutions, and require no or only subtle modification of the investigated molecules. Microfluidics-based methods are well suited for such tasks, and in this paper, we describe an experimental setup meeting the criteria above.

### Basic principles of the experimental method

The inspiration for this method was published by Arosio et al., where green fluorescent polystyrene particles, α-synuclein molecules, antibody fragments, and other nanoparticles were injected into a microfluidic device in order to measure their lateral movement via diffusion, based on which their hydrodynamic radii could be approximated and compared to *a priori* data generated with dynamic light scattering (DLS). [3] This technique was also explored by Gang et al. with polydisperse mixtures including up to three components with distinct sizes: α-synuclein fibrils, small unilamellar vesicles (SUVs), and SUVs with α-synuclein fibrils bound to the external surface of their membranes. [4] Adapting this technique to PSD proteins and their complexes where particles have a dynamic range from nm to μm proved challenging but the theory behind it is very similar. In solutions unrestricted particles are in perpetual motion via diffusion. The smaller they are the faster they move, as described by the Stokes-Einstein equation:

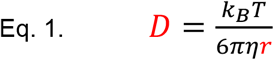

where D is the diffusion coefficient of the particle, k_B_ is the Boltzmann constant, T is the absolute temperature, η is the viscosity of the (liquid) medium, and r is the hydrodynamic radius of the (spherical) particle. It is important to note that this equation only works in case of globular particles and a liquid with low Reynolds number, which is characteristic of laminar flow. So, the method requires a device that achieves laminar flow for protein samples, from now on referred to as analytes. Additionally, the device has to arrange particles into a predetermined position from where they can freely diffuse during the laminar flow. A microfluidic focuser with the appropriate layout can achieve these criteria by compressing the analyte into the middle of a channel with buffer streams from both sides.

This method is also based on the assumption that diffusing particles would approximate normal distribution. Since their motion is primarily governed by diffusion in directions that are perpendicular to the laminar flow’s direction, it is logical to examine it in one such direction. Thus, the distribution of particles can be described through Brownian motion with one spatial dimension:

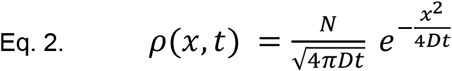

where t is the time elapsed since diffusion began, N is the number of particles that start from the origin at time t = 0, x is the distance from the origin, and ρ is the particle density at distance x at time t. Concurrently, the mathematical description of a Gaussian function is as follow:

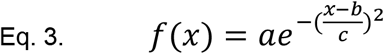

where a is the amplitude, b is the center and c is the standard deviation of the bell curve, while f(x) is the value of the function at position x. If the time component in Eq. 2. is set to a constant value, then Eq. 2-3. match as

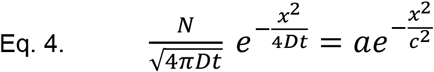

if the Gaussian function is centered on the origin (b = 0). The following expressions can be derived from Eq. 4.:

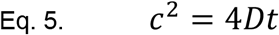

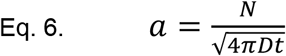

From Eq. 5-6. the diffusion coefficient D and the number of particles N can be written as follows:

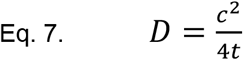

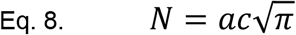

And so, the number of particles can be approximated if the amplitude and standard deviation of the Gaussian function describing their distribution at a given point are known. More importantly, their diffusion coefficient, and thus their size, can be approximated as the incline of the linear function fitted to the points given by the Gaussian functions’ variances and the associated time components (Fig. 1.).

**Fig. 1.**
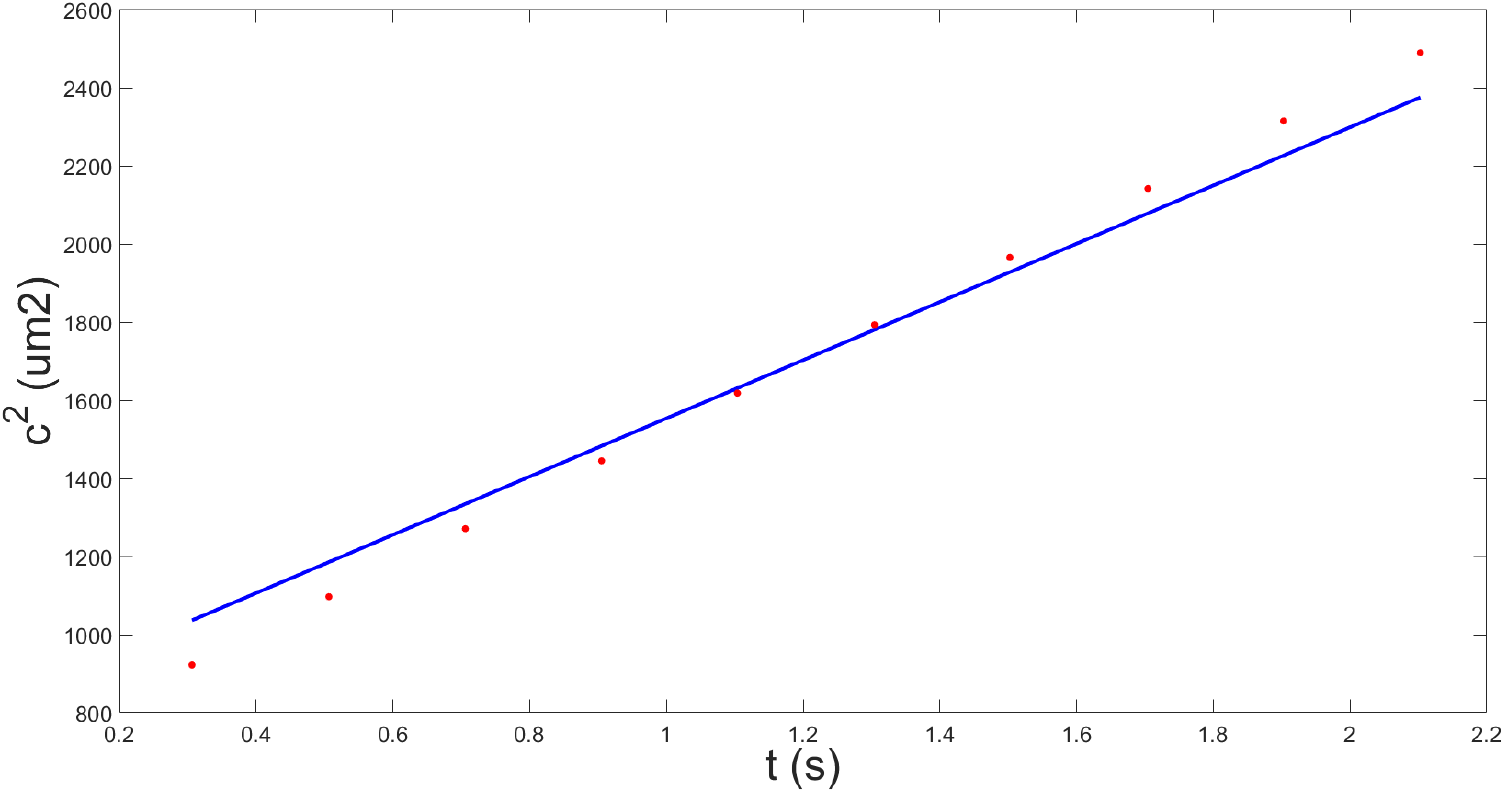
The variance of Gaussian functions (c^2^) fitted to fluorescent intensity profiles measured at different distances from the front of the channel (where the analyte was focused). The x-axis shows the average time (t) it takes for the particles to reach a given measurement point at a constant flow rate (see Measurement protocol). The analyte was Enhanced Green Fluorescent Protein (EGFP).

This requires multiple measurements at multiple points in time during the laminar flow, which can be achieved by setting up multiple measurement points along a straight channel added to the microfluidic focuser.

### Postsynaptic densities

PSDs are multilayered cellular components situated on the internal surface of postsynaptic membranes. These disk-shaped compartments are rich in proteins and nucleic acids, including RNA-binding proteins, actins, membrane-associated guanylate kinases (MAGUKs), and scaffolds. Their primary constituents are the postsynaptic density protein 95 (PSD-95), the guanylate kinase-associated protein (GKAP), the synaptic Ras GTPase-activating protein 1 (SynGAP), as well as various Shank and Homer proteins. [5-10] PSDs are also associated with transmembrane proteins, such as the N-methyl D-aspartate receptor (NMDAR). [11] They are the primary cellular elements that process incoming synaptic transmissions, and their structural changes exhibit a strong correlation to synaptic strength and plasticity, which are pillars of higher biological functions such as memory and learning. [12] Thus the phase separation of PSD proteins on the molecular level, synaptic plasticity on the cellular level, and memory on the systemic level are all connected to each other. The functional importance of PSDs is further emphasized by studies of the wake/sleep cycle suggesting that synaptic strength is renormalized during sleep. [13]

The structural organization of PSDs is governed by various complex biochemical processes, including liquid-liquid phase separation (LLPS). Protein phase separation is a complex biochemical process, typically initiated by multivalent interactions between multi-domain proteins and RNAs. The phenomenon produces so called membraneless organelles (MLOs) that often play the role of dynamically changing storage units for the components of certain cellular processes, such as RNA transcription, chromatin regulation, or the organization of PSDs. [12, 14, 15] LLPS is a specific type of protein phase separation characterized by the formation of distinct phases where certain solutes appear in high concentrations, resulting in liquid-like properties. This particular type of the phenomenon is driven by multivalent proteins appropriately referred to as “drivers.” [16] The role of LLPS in structural organization is indicated by an experimental model constructed from two abundant components of PSDs, SynGAP and PSD-95. This simple model has been observed to self-organize into highly condensed, PSD-like droplets *in vitro*. [17] The importance of phase separation in the organization of PSDs has since been reinforced by more complex in vitro models, one including GKAP, Shank3, and Homer3, in addition to SynGAP and PSD-95. [18]

The diffusion-based analytic method presented here was tested on a two-component system from the network above. One of the components is GKAP, a known scaffolding protein that participates in the regulation of NMDA receptors. It contains a high ratio of disordered regions and multiple binding sites, two of which had been identified as LC8 binding motifs. [19] The other component is the dynein light chain LC8 protein that can form multivalent interactions with various intrinsically disordered regions and proteins (IDRs / IDPs). [20] It is also known to form dimers that can bind two additional ligands, suggesting potential for the initiation of MLO-formation through multivalent interactions. This potential is further supported by the level of disorder within GKAP and the binding partners of LC8, since IDPs are generally prone towards phase separation, as observed in ca0se of the RNA-binding protein Fused in sarcoma (FUS), the TAR DNA-binding protein 43 (TDP-43), and the intrinsically disordered intracellular domain of Nephrin (NICD). [21-23] GKAP and LC8 are also known to form hetero-oligomeric complexes, the exact stoichiometry of which has only been explored recently. [24-25] Based on these developments of our understanding of GKAP-LC8 complexes, the two-component system constructed for testing the diffusion-based analytic method consisted of LC8 and GKAP molecules with a 4:2 stoichiometry (2:2 stoichiometry of LC8 dimers and GKAP monomers). The GKAP construct containing both LC8-binding regions is referred to as GKAP-DLC2. GKAP’S PDZ-binding motif (its construct referred to as GKAP-PBM) was also investigated, as a standalone particle at this point. Apart from GKAP and LC8, the protein Drebrin has also been analyzed via this method, though only as a standalone particle, as it does not exhibit direct interactions with GKAP or LC8, but the EVH1 domain of Homer, another component of the PSD. The Drebrin construct used in this study is referred to as D233.

## Results

### Development of the microfluidic device

Developing microfluidic devices was and still is a process both collaborative and iterative. First, an initial layout is agreed upon, and then a couple of devices are manufactured for testing and evaluation, after which the layout gets reworked based on the observations. All layouts were designed in Autodesk AutoCAD 2025.

It was quickly realized that the angle of the intersection was directly proportional to the probability of backflow, where the two side streams would flow towards the middle inlet as well as the outlet, halting the middle stream that must be set to a lower flow rate (see the “Measurement protocol” section). Although decreasing the angle from 90° to 60° significantly reduced the likelihood of backflow, lowering it further to 45° did not seem to make more of a difference. Since the inlets cannot be placed any closer to the edge of the device, the intersection has to be placed further away to achieve lower angles, which leaves less space for the main channel where diffusion is to be observed. Because of these reasons, future layouts all incorporated intersections where the inlets meet at an angle, but that angle would be at least 60° to conserve space for the main channel.

Another way of reducing the likelihood of backflow is introducing a resistance to each inlet. However, these components require space, pushing the intersection further away, leaving less space for the main channel. A compromise of smaller resistances on the side inlets and a 67° angle at the intersection has proven to reduce the probability of backflow to a negligible level, while conserving as much space for the main channel as possible. In addition to these changes, the main channel has been widened to reduce resistance towards the outlet. An additional benefit of this was that it gives more space for diffusion before particles reach the side walls of the channel. This allowed for lower flow rates, which gave larger particles more time for diffusion. The main channel was also elongated by adding turns to it, in order to give particles even more time for diffusion before they reach the end of the channel. However, each of these turns introduced a centripetal force to the particles that is perpendicular to the flow, undermining the basic principle that only diffusion should affect particles in that direction.

Another way of elongating the main channel is increasing the size of the device. At first, this possibility was avoided because larger devices cannot be bonded to cover plates, only glass slides that are thicker, preventing the usage of high resolution lenses with 60x magnification. However, as the measurement protocol evolved, the usage of a 60x lens became unnecessary regardless of the microfluidic device. It was also established that it is possible to produce high quality images with devices bonded to glass slides.

A device consisting of three design units is the largest size that still fits on a single glass slide, therefore the 54 mm long main channel that the above layout contains is the longest channel possible without including turns. The layout was also improved with markers at eleven equidistant measurement points along the main channel, the first of which is 2.5 mm after the intersection. These measurement points help tracking the exact locations of recorded images, which is required to determine the distance between recorded profiles, which in turn is vital for determining the average time particles spend between them (see Fig. 2.).

**Fig. 2.**
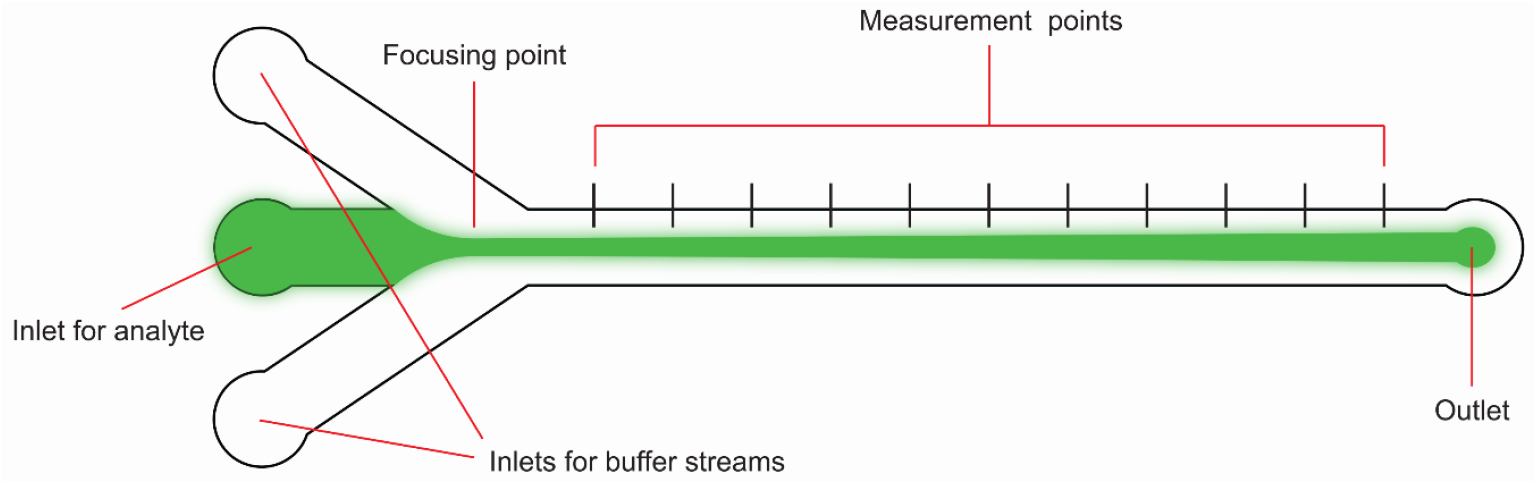
Basic layout of the microfluidic focuser that has a channel attached to it with multiple measurement points.

**Fig. 3.**
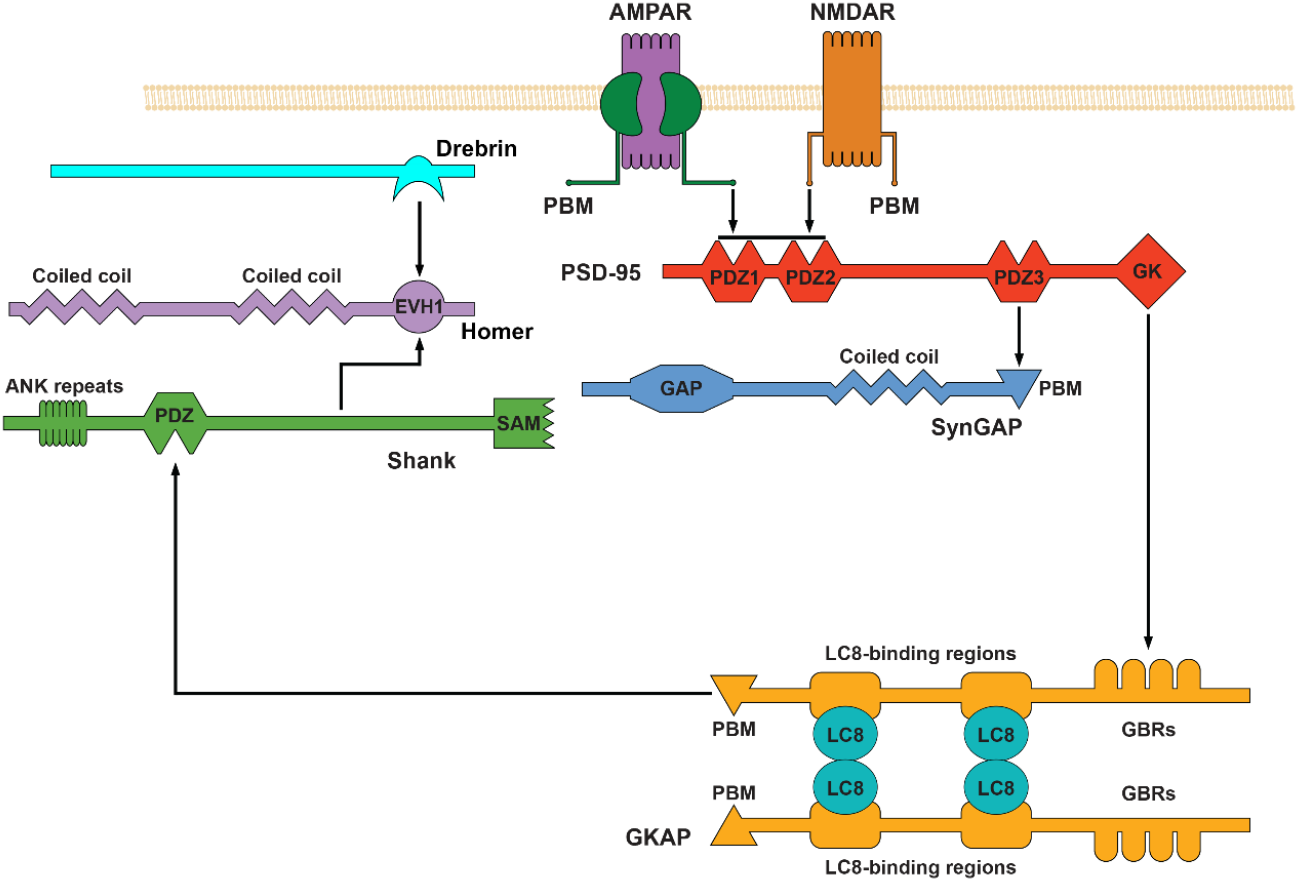
Schematic illustration for an *in vitro* PSD model. Proteins are depicted in arbitrary orientations, serving a simpler representation of the selected interactions. This study primarily focuses on GKAP’s interactions within this network, specifically its PDZ-binding motif and the hexameric complex formed with LC8 dimers that bind to GKAP molecules at multiple positions.

**Fig. 4.**
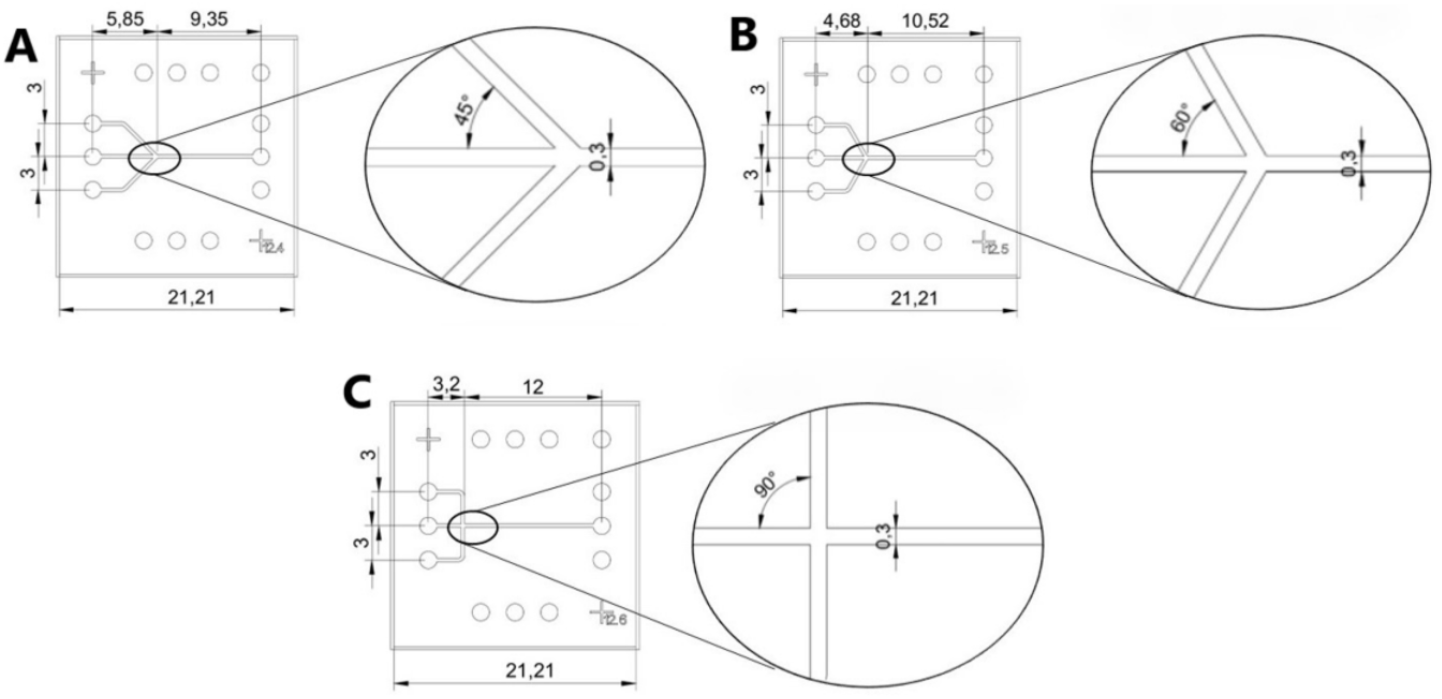
Layout of three early designs that were simple microfluidic focusers with three inlets, one outlet. Flow quality was tested with different intersections at angles of 45° (A), 60° (B), and 90° (C). All parameters are in mm, the channel height is 20 μm. The width of the main channel and the individual inlets were the same, and the entire device was one cell long in all three cases.

**Fig. 5.**
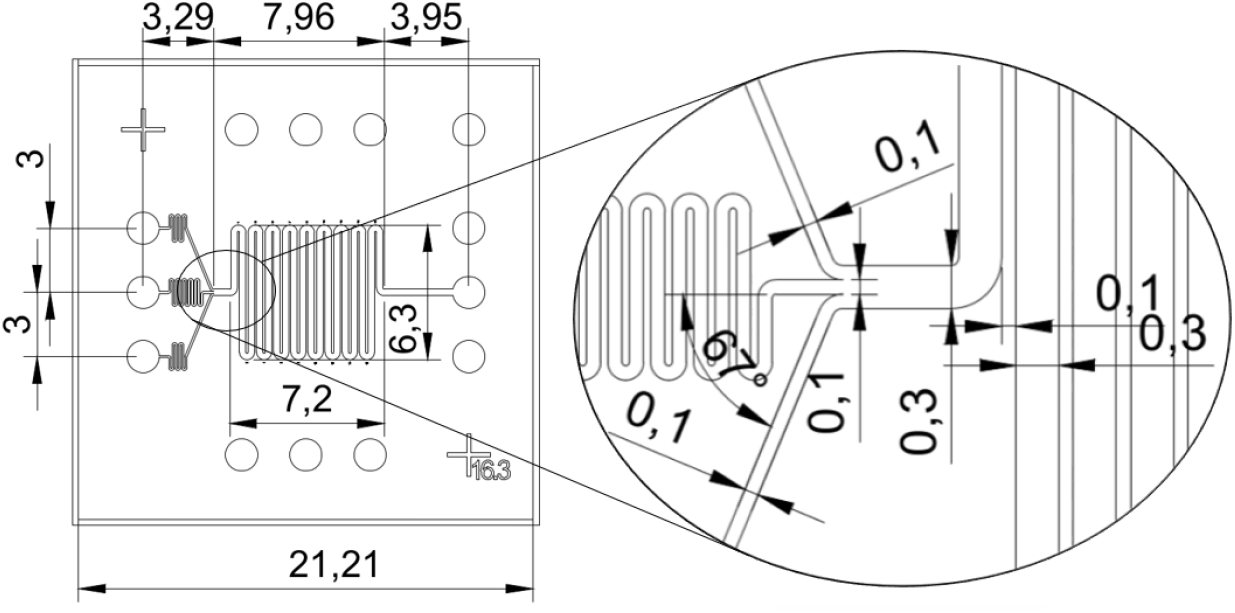
Layout of an improved design where the inlets meet at a 67° angle. All parameters are in mm, the channel height is 20 μm.

**Fig. 6.**
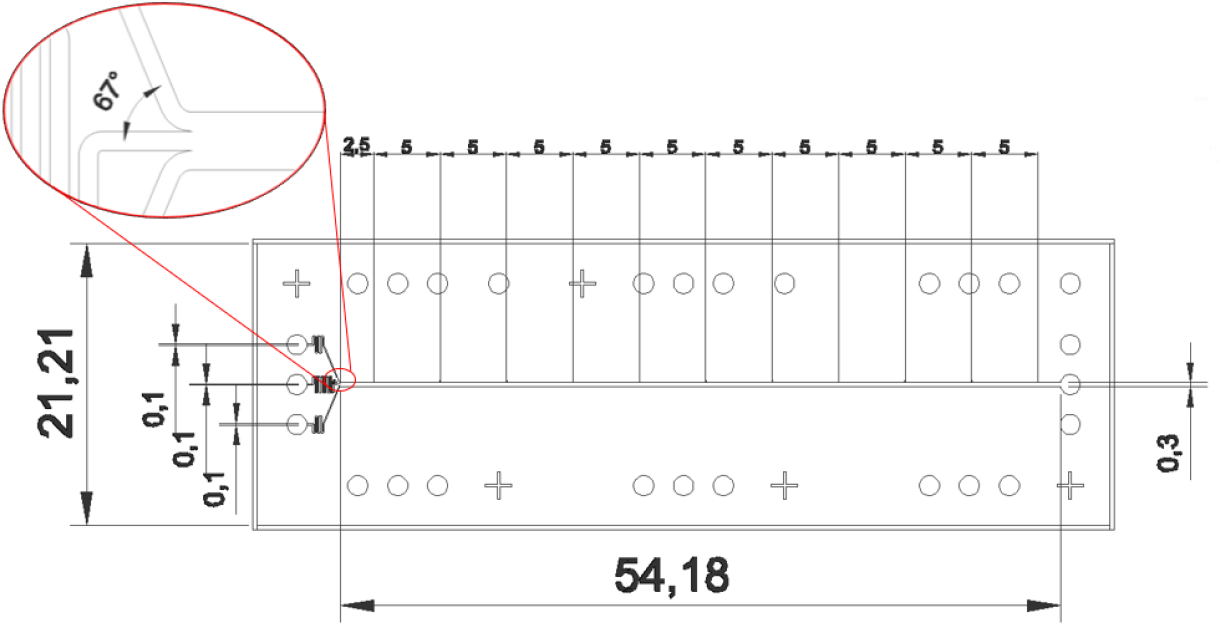
Layout of the first device that consisted of two design units. All parameters are in mm, the channel height is 20 μm.

**Fig. 7.**
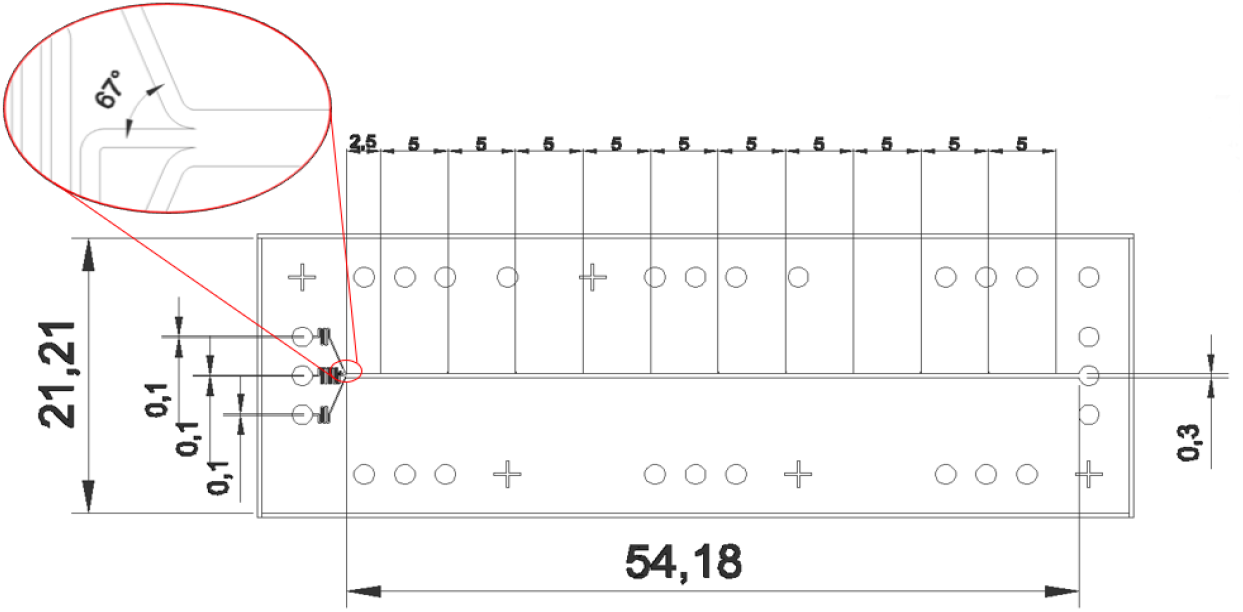
Layout of the current device. The length of the device was extended to three cells. All parameters are in mm, the channel height is 20 μm.

**Fig. 8.**
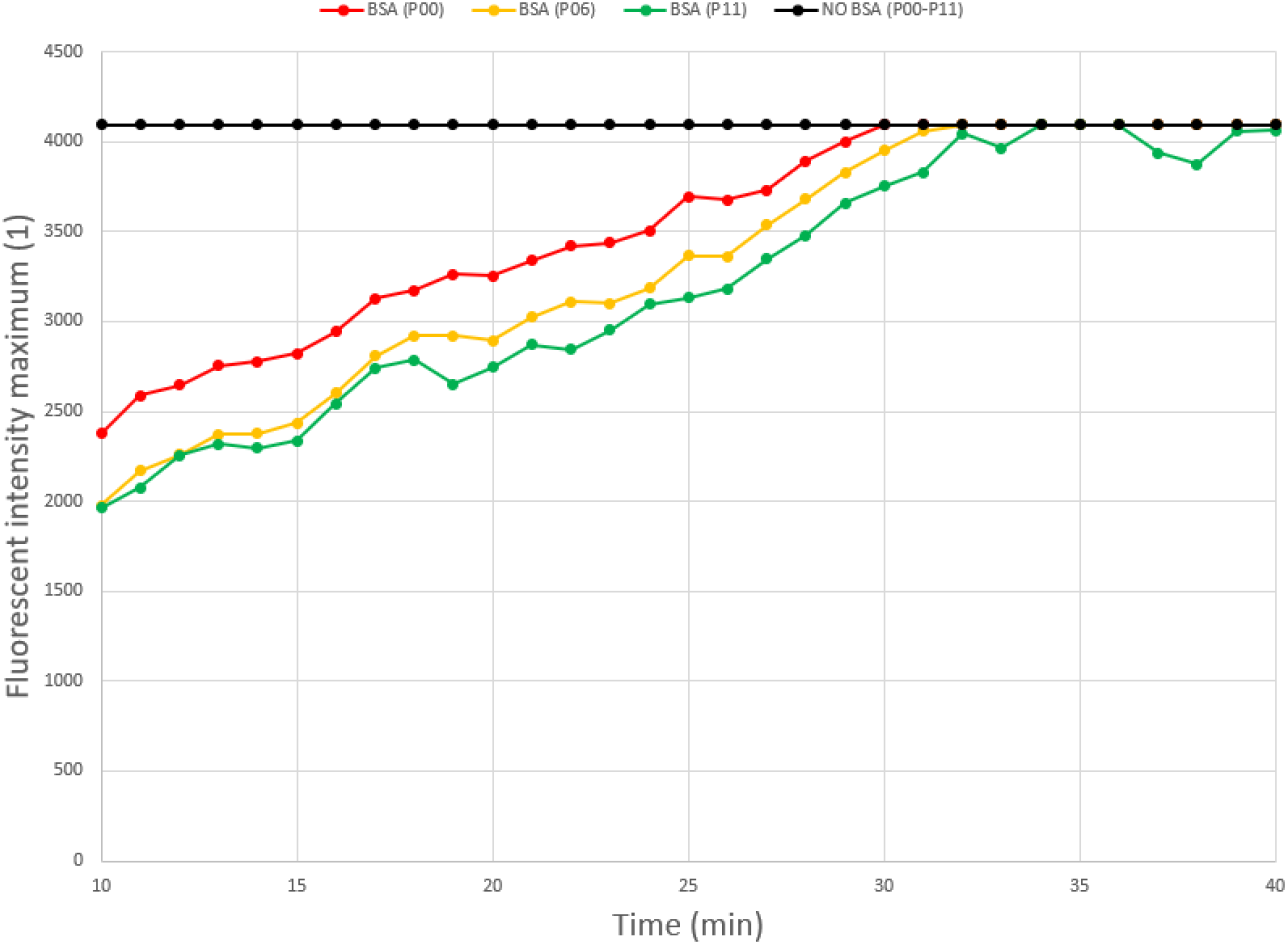
Maxima of fluorescent intensity profiles at the beginning of the main channel (P00), at the middle of the channel (P06), and at the end (P11), measured every minute for half an hour with and without treating the microfluidic device with BSA. In each case, the analyte consisted of fluorescent microspheres with a nominal diameter of 0.05 μm, and the first measurement took place 10 mins after the analyte had reached the beginning of the channel (see Measurement protocol).

**Fig. 9.**
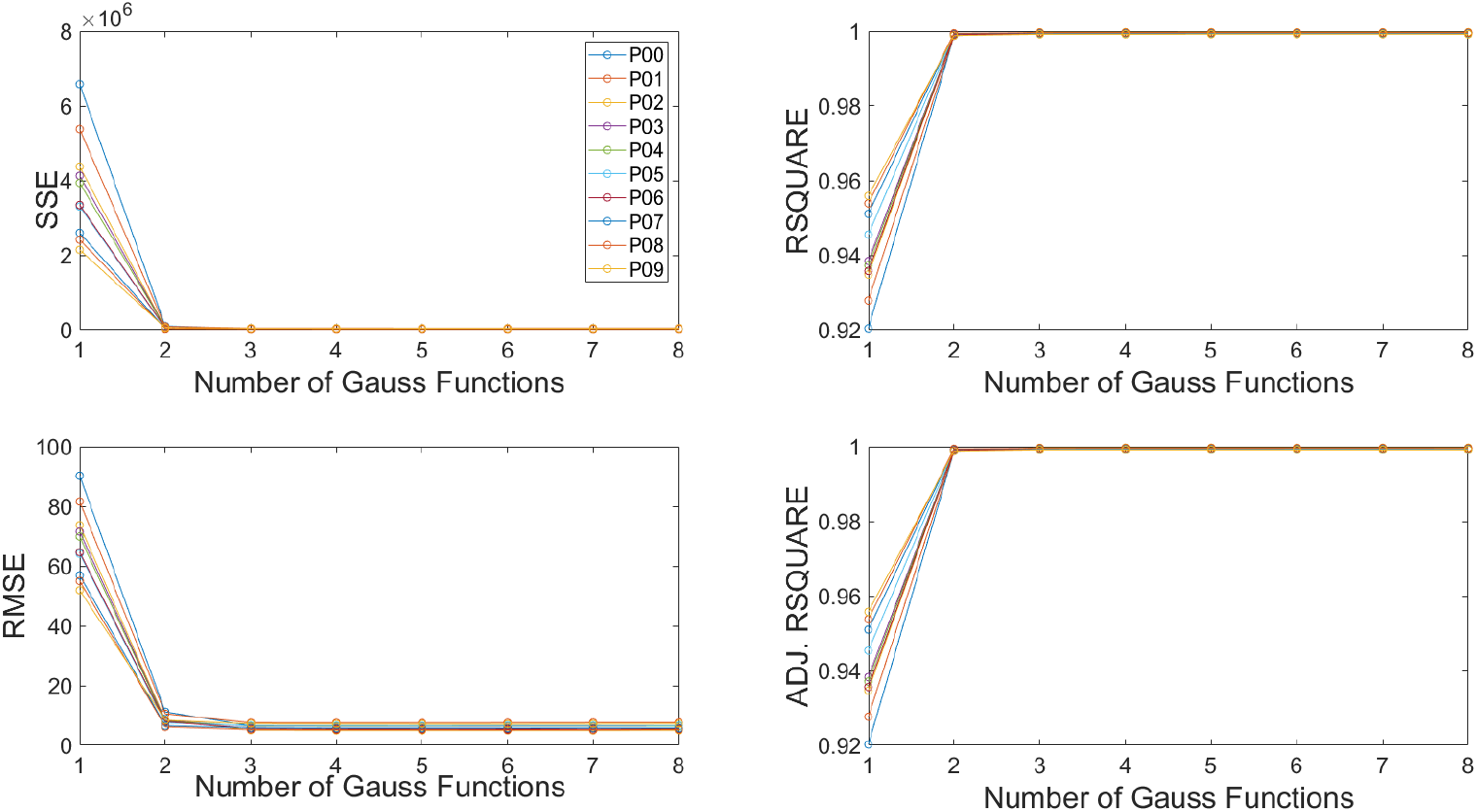
Goodness of fit (GoF) metrics, calculated for all measurement points (P00-P11) and eight levels of fit complexity (1-8). The first level is fitting a single Gaussian function to the measured data at each measurement point, while higher levels mean fitting the linear combination of multiple (2-8) Gaussian functions. The analyte was EGFP for this particular measurement, but all analytes (see Table 1.) exhibited a similar trend in all GoF metrics.

**Fig. 10.**
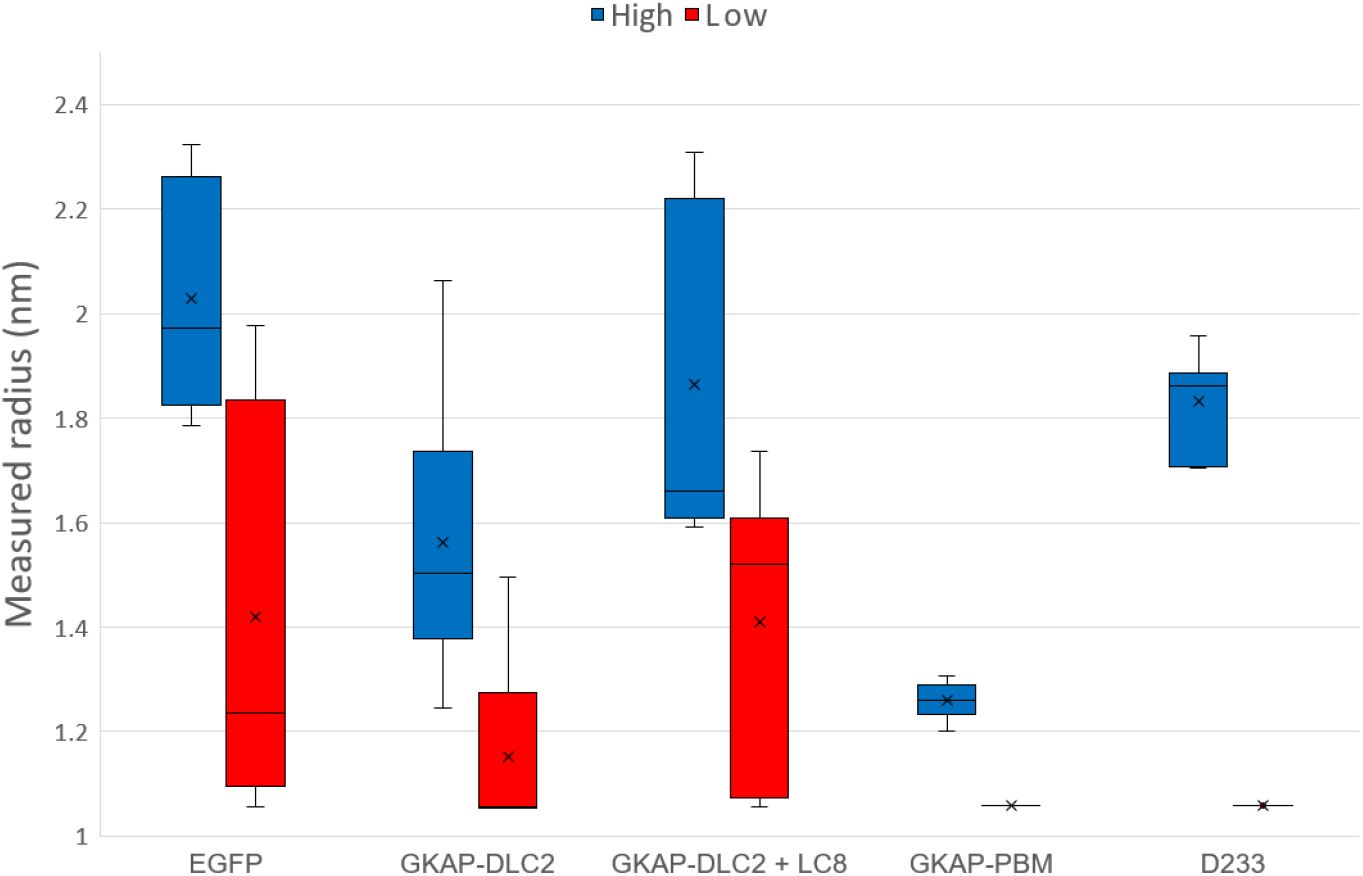
Box plot of approximated radii for different particle types. The minimum boundary for approximate radii was set to 1 nm in the analytic software. Due to rounding errors in the various operations executed on the measured data (see The analytic software package), the actual lower limit is about 1.05 nm, which is reflected by the boxes of low values.

**Fig. 11.**
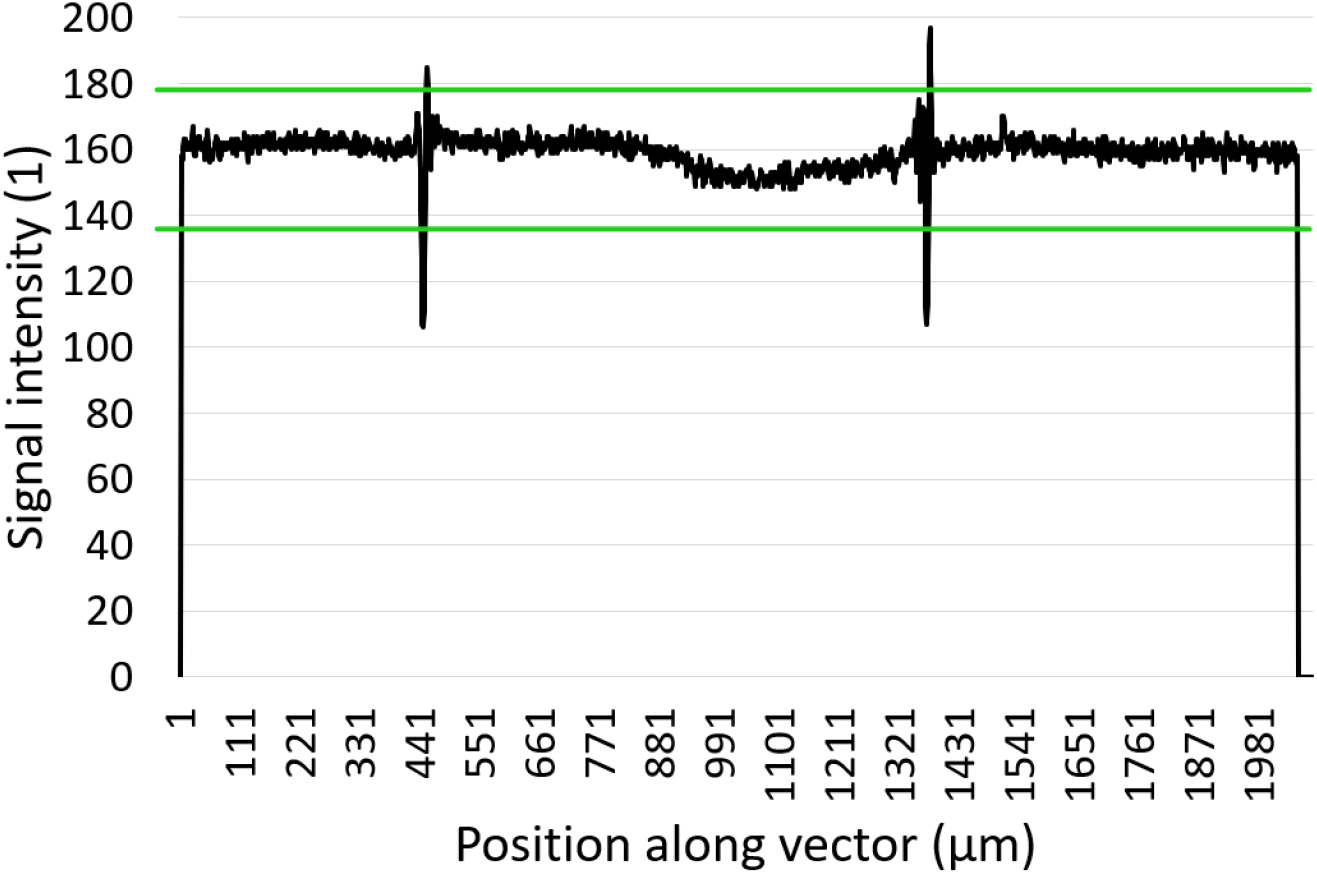
Brightfield intensity profile of a FITC+GKAP-DLC2 complex sample (**black**) with horizontal lines (green) at its average minus its standard deviation (135.66) and at its average plus its standard deviation (177.53). The intensity drops to zero towards both ends where the vector includes points outside the image. The two large spikes at 140-160 μm and 340-360 μm are the sidewalls of the channel that become relevant in the normalization process (see below).

**Fig. 12.**
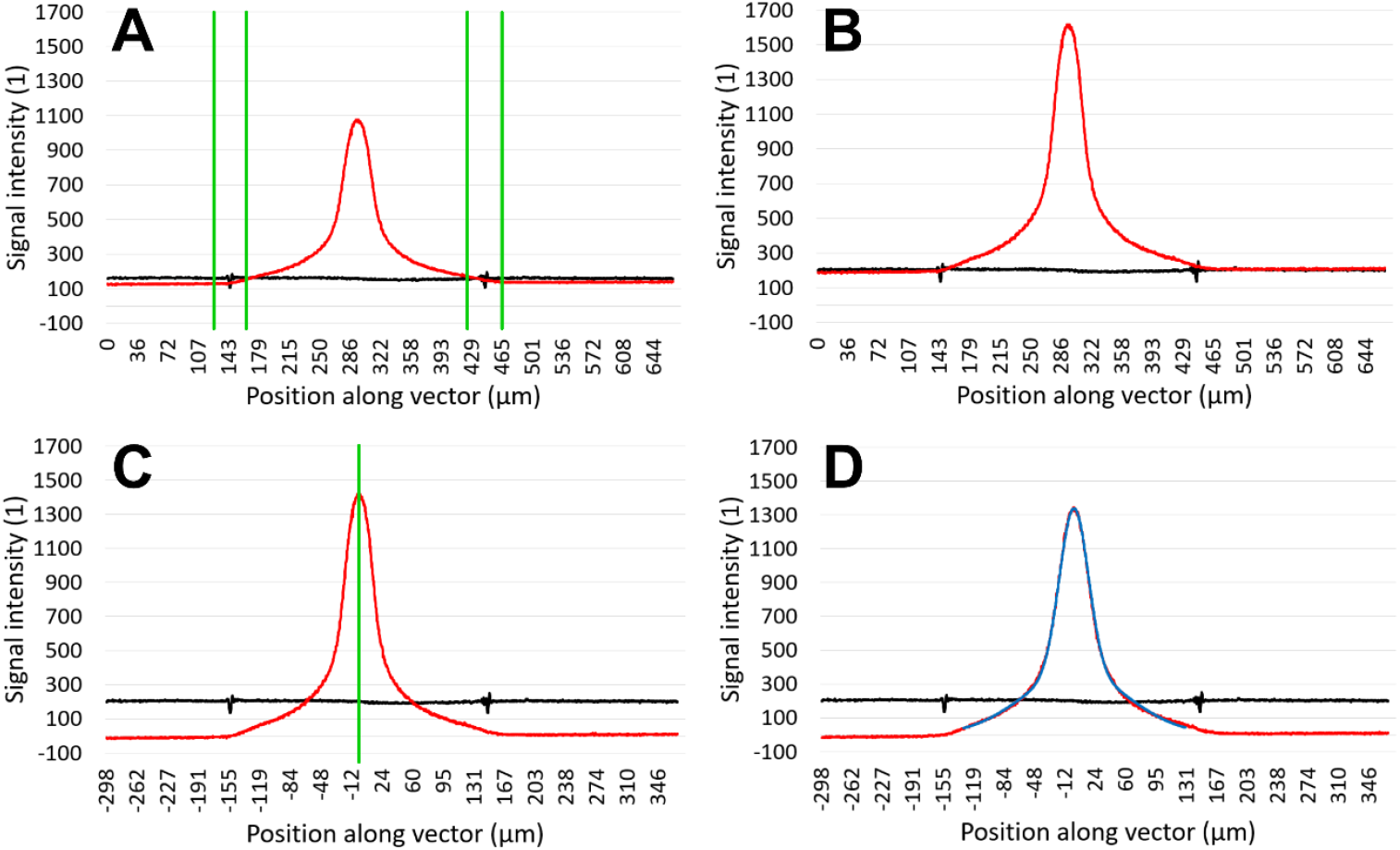
Four major steps from cropping the brightfield (**black**) and fluorescent (red) profiles to fitting Gaussian functions. **A)** Sections around the spikes in the brightfield profile reveal the position of the channel’s sidewalls (green). The thickness of the sidewalls is doubled to account for the uncertainty in the relation between the spikes’ position and the middle of the actual sidewall. **B)** Both brightfield and fluorescent profiles are multiplied by their respective coefficients to move their baseline to the standard 200. **C)** The fluorescent profile is further normalized according to its integral, then aligned with the (Signal intensity = 0) axis as well as centered around its peak value (green). **D)** A linear combination of two Gaussian functions (blue) is fitted to the fluorescent profile inside the channel.

### Size approximation

Four GoF metrics were considered when evaluating the different models: Sum of squares due to error (SSE), root mean squared error (RMSE), R-square, and degrees of freedom adjusted R-square. Since all analytes were assumed to be monodisperse, it was favorable to choose a model with low complexity. Therefore, measured data were fitted with the linear combination of two Gaussian functions, as goodness of fit metrics do not improve significantly at higher levels of complexity.

**Table 1.** List of analytes, complete with their measured radii via DLS and the diffusion-based approach. In the case of GKAP-PBM and D233 the DLS measurements were replaced by approximations based on their molecular weights, respectively. Each set of DLS measurements yielded a mean hydrodynamic radius and an associated standard deviation. Each set is listed individually where applicable. The mean values and standard deviations for the diffusion-based approach (approximate radii) were calculated from multiple measurements. EGFP samples were used for the calibration of the experimental setup. PSD protein samples included D233, GKAP-PBM, and GKAP-DLC2 molecules, as well as GKAP-DLC2+LC8 hexamers, respectively. Drebrin and GKAP molecules were labelled with fluorescein (FITC). Since the analytic software was set to fit the linear combination of two Gaussian functions to the measured data (see Fig. 9.), the output from each measurement was a pair of approximate radii. Column three includes both values, while columns four and five only include either the higher or the lower value of the same output.

| Analyte | Expected radius (nm) | Approximate radius (nm) | Higher approx. radius (nm) | Lower approx. radius (nm) |
| --- | --- | --- | --- | --- |
| GKAP-PBM | 1.39 | $1.16 \pm 0.10$ | $1.26 \pm 0.03$ | $1.06 \pm 0.00$ |
| GKAP-DLC2 | $1.27 \pm 0.72$<br>$1.90 \pm 1.00$ | $1.36 \pm 0.30$ | $1.56 \pm 0.24$ | $1.15 \pm 0.18$ |
| GKAP-DLC2 + LC8 | $2.17 \pm 0.44$<br>$3.11 \pm 0.76$ | $1.64 \pm 0.36$ | $1.91 \pm 0.30$ | $1.38 \pm 0.27$ |
| D233 | 1.81 | $1.45 \pm 0.39$ | $1.83 \pm 0.09$ | $1.06 \pm 0.00$ |
| EGFP | $2.72 \pm 1.01$ | $1.72 \pm 0.42$ | $2.03 \pm 0.20$ | $1.42 \pm 0.36$ |

The approximate radii are quite close to their expected values for such small particles. However, since the diffusion of smaller particles is much more significant at the same flow rate, the method might be biased towards them when fitting Gaussian functions to the measured profiles. This is somewhat mitigated by discarding the lower value from the output. Both DLS and the diffusion-based approach reflect that GKAP-DLC2+LC8 hexamers are larger than GKAP-DLC2 monomers, which are in turn larger than GKAP-PBM domains. However, in the case of DLS, it is unclear whether EGFP molecules are larger than GKAP-DLC2+LC8 hexamers. In all cases, the standard deviation is lower for the diffusion-based approach, highlighting that at the ends of the 1 nm - 10 μm range of DLS (Zetasizer Ultra/Pro), its precision becomes increasingly system-dependent.

## Discussion

Measuring the size distribution of particles within a flowing solution via microfluidics and fluorescent microscopy is a working concept that has limitations in distinguishing particles the sizes of which are of the same magnitude. The method is biased towards smaller particles present in the solution, and the results yielded by it must be cross-referenced with that of samples with known particle sizes. The method provides accurate approximations for particles below 10 nm in diameter. It becomes increasingly inaccurate for larger particles, yielding completely wrong approximations for particles in the 1 μm range and above (see Supplementary).

The length of the PTFE tubing connected to the microfluidic device’s inlets could be further decreased with the fabrication of specialized equipment that holds the syringe pumps closer to the device and mimics the motions of the stage without interfering with the microscope. This would reduce the time it takes for the solutions to reach the device as well as the minimum volume of solutions required for a measurement.

Ultimately, the method in its current state cannot reliably distinguish the observed proteins from their complexes, but it is sufficient for distinguishing proteins and their complexes from larger particles. The accurate measurement of larger particles would require a more refined model that describes signal forms with polynomial functions, creating a library from particles of known sizes to which new data could be compared to. This improved method would still be limited, however, to the measurement of particles that conform to already acquired data that can only cover a set of distinct sizes.

## Methods

### Device fabrication

The molds for microfluidic devices were fabricated using a soft lithography and polydimethylsiloxane (PDMS) replica molding techniques. In this process, a negative photoresist height of 20 μm (SU-8 2015, Microchem Corp., Newton MA, USA) is applied to the top of a silicon wafer by spin-coating (WS-650-23, Laurell Technologies Corp., Lansdale PA, USA). Channel design was applied on the surface by laser writing (μPG 101, Heidelberg Instruments Mikrotechnik GmbH, Heidelberg, Germany). After the development of the mold the PDMS (SYLGARD 184 Silicone Elastomer Kit, Dow Corning, Midland, MI) base and curing agent were mixed in a 10:1 ratio, degassed, poured over the mold, and cured at 70°C for 90 min. Once the polymerization process is complete, the PDMS was removed from the mold surface, the inlets and outlets were processed. The PDMS slice containing the microfluidic channel was bonded to a glass slide using plasma treatment (Zepto, Diener electronic GmbH, Ebhausen, Germany).

PTFE tubing (Masterflex Microbore Transfer Tubing MFLX06417-11, Cole-Parmer Instrument Company, LLC, Vernon Hills IL, USA), was inserted into the device inlets and outlets, then connected to 2 ml syringes and NE-1002X syringe pumps via a 27 Gauge needle tip (27 Gauge Clear Precision Tip 0.5” Long, Adhesive Dispensing, Ltd, UK). The PTFE tube on the outlet is approx. 15 cm long and leads into a beaker that is also positioned on the stage of the microscope. The PTFE tubes on the inlets are approx. 25 cm in length, which is the minimum length required to keep the tubing from becoming taut as the stage moves the device around, minimizing the risk of it slipping out of the inlets.

### Protein expression and purification

The GKAP-DLC2 construct was designed to include both LC8-binding motifs of GKAP with extended flanking regions (10 residues on the N-terminus, 14 residues on the C-terminus). The segment spanning residues 655-711 in the *Rattus norvegicus* GKAP isoform 3 “GKAP1a” (UniProt ID: P97836-5) was selected. The GKAP-PBM construct incorporated the PDZ-binding motif EAQTRL and its extensive flanking region of 37 additional residues. Both inserts were cloned to an altered pEV vector (Novagen) that contains an N-terminal 6xHis tag and a tobacco etch virus (TEV) protease cleavage site. The actual constructs contained four extra residues (GSHM) at the N-terminus, remaining from the expression tag. The *Rattus norvegicus* DYNLL2 gene (UniProt ID: Q78P75) in the pEV plasmid vector is 100% identical to the human ortholog (UniProt ID: Q96FJ2). The pEV vector also contains an N-terminal 6xHis tag, the TEV protease cleavage site and four residues (GSHM) at the N-terminus.

All three protein constructs were produced in BL21 (DE3) *E. col*i (Novagen) cells, transformed with the vectors, grown in LB media, induced with 1 mM IPTG (Isopropyl β-D-1-thiogalactopyranoside) at 6 MFU cell density, and the recombinant proteins were expressed at 20°C overnight. The centrifuged cell pellets were stored at -20°C for further usage. Cell pellets were lysed by ultrasonic homogenization in 10% cell suspension using a lysis buffer (50 mM NaPi, 300 mM NaCl, pH 7.4). In the case of GKAP-DLC2, denaturing-renaturing IMAC purification was applied (denatured with 6 M GdnHCl, 50 mM NaPi, added to 5 ml Nuvia™ Ni-affinity column (Bio-Rad), then bound proteins were renatured with native buffer (50 mM NaPi, 20 mM NaCl, pH 7.4). After washing, elution was performed with 250 mM imidazole and was followed by His-tag removal with TEV protease. For LC8 and GKAP-PBM, the same purification protocol was used but after ultrasonic homogenization and centrifugation, the supernatant was immediately purified with IMAC Nuvia Ni-affinity column without denaturation-renaturation.

Protein samples were concentrated by ultrafiltration using Amicon® Ultra Centrifugal Filter with 3 kDa molecular weight cut off value, and the buffer was changed to low salt NaPi Buffer (50 mM NaPi, 20 mM NaCl, pH 6.0). Proteins, except GKAP-PBM, were further purified by ion exchange chromatography (IEC), using 5 ml High Q column with the same buffer (50 mM NaPi, 20 mM NaCl, pH 6.0). Recombinant proteins were collected in the flow through fraction. After another step of protein concentration, proteins were further purified with size exclusion chromatography (SEC) on a Superdex™ 75 Increase 10/300 GL 24 ml column, the buffer was 50 mM NaPi, 20 mM NaCl, pH 6.0. Later 5 mM TCEP (pH adjusted to 7.4 with NaOH) was added to GKAP-DLC2 and LC8 samples individually. The concentration of LC8 and GKAP-PBM was measured by its absorbance at 280 nm using a NanoDrop2000 photometer, while the concentration of GKAP-DLC2+LC8 hexamers was measured with Qubit Protein assay. The molecular weight of GKAP-DLC2, LC8, and GKAP-PBM monomers was determined to be 7.01 kDa, 10.6 kDa, and 5.2 kDa, respectively, validated via SDS-PAGE.

Drebrin (D233) was produced by using the full sequence in the *Homo sapiens* isoform Q16643 as template for cloning the segment 233-317 into NdeI and HindIII sites of a modified pET-15b vector along with an N-terminal 6xHis-tag and a TEV cleavage site (ENLYFQG). This construct was also produced in BL21 (DE3) *E. coli* cells and grown in LB media. After inducing with 1 mM IPTG, cells were incubated for 3 h at 37 °C before harvesting by centrifugation. The lysis buffer (50 mM NaPi, 300 mM NaCl, 5 mM β-mercaptoethanol, pH 7.4) also contained 1 mM AEBSF protease inhibitor cocktail (Thermo Scientific AEBSF Protease Inhibitor #78431). The same buffer, without the protease inhibitor cocktail, was added to the Ni-affinity column while applying IMAC purification to D233. Elution was performed with 500 mM imidazole, and the His-tag was cleaved with TEV protease. The sample was further purified via SEC on a Superdex™ 75 Increase 10/300 GL 24 ml column equilibrated with buffer suitable for FITC labelling (50 mM NaPi, 20 mM NaCl, pH 8.0). The molecular weight of D233 was determined to be 10.32 kDa, reinforced by SDS-PAGE.

### Preparation of samples

Protein samples were labelled with the Green-fluorescent Fluorescein-EX (FITC) labelling kit (ThermoFisher cat. num. F10240) using the following protocol: The buffer of GKAP-DLC2, GKAP-PBM, and D233 was changed to 50 mM NaPi, 20 mM NaCl, pH 8.0 as suggested by the manufacturer. Based on absorbance measurement with Nanodrop, the concentration was 3.5 mg/ml and 2 mg/ml respectively. 0.5 mL and 1.5 mL samples were added to one vial containing the reactive dye, respectively. After one hour of incubation and stirring at room temperature, the vials were stored at 4°C overnight (for 16 hours). The preparation of Drebrin samples slightly diverged here, only being incubated for 1 hour. After the labeling reaction, any unbound reactive dye was separated from the labeled protein with size exclusion chromatography (SEC). 0.5 mL samples were injected one by one to a Superdex™ 75 Increase 10/300 GL 24 ml column, the flow speed was between 0.8 ml/min (with the same buffer as used for the labeling: 50 mM NaPi, 20 mM NaCl, pH 8.0). Unlabeled LC8 dimers were added to the fluorescein-labeled GKAP-DLC2 with 2:2 stoichiometry (i.e. 2 LC8 dimers to 2 GKAP-DLC2 monomers). The final volume of labelled protein solutions had to be at least 300 μl per measurement. All protein samples were focused in the microfluidic device with low salt NaPi buffer streams (50 mM NaPi, 20 mM NaCl, pH 8.0).

Additionally, Enhanced Green Fluorescent Protein was used for testing the various iterations of the microfluidic device. EGFP analytes were focused with a PBS buffer (137 mM NaCl, 2.7 mM KCl, 10 mM Na_2_HPO_4_, and 1.8 mM KH_2_PO_4_, pH 7.4). The analytes were prepared by diluting 2 mg/ml EGFP stock solutions with an equal amount of PBS buffer. The 1% BSA solution used for the surface treatment of microfluidic devices was prepared in stock by dissolving 0.05 g BSA powder (VWR Chemicals Bovine Serum Albumin, cat. no. 97061-420) in 50 ml of PBS buffer, then aliquoted to 1 ml Eppendorf tubes to be frozen for later use.

### Measurement protocol

A measurement requires a microfluidic device connected to three syringe pumps. The device is placed into a Nikon Ti-2 E inverted microscope with a motorized stage and filter turret that detects signals passing through a FITC filter (Excitation: 480/30, Dichroic mirror: 505, Barrier filter: 515) and recorded with an Andor Zyla 4.2 camera. A Nikon CFI Plan Apochromat Lambda D 20X lens is used for measurements, because it is not possible to record the entire 300 μm width of the main channel at higher magnifications. The device is also taped to the stage to minimize the risk of even the slightest of movements as the stage repeatedly repositions it during the measurement.

The internal surface of the device is treated with BSA prior to the measurement in order to slow down the process of fluorescent particles sticking to the surface. This is achieved by filling each syringe with at least 600 μl of 1% BSA solution that is pumped into the device at a flow rate of 20 μl/min. After the solutions from all three syringes have reached the device the aforementioned flow rate is maintained for an additional 15 minutes to properly degas the entire system. This is followed by decreasing flow rates to 4 μl/min, which is maintained for another 10-15 minutes, since the flow rate of the solution exhibits a hyperbolic decline after lowering its value on the syringe pumps. While the flow is slowing down the markers of the predefined measurement points are manually brought into focus one by one, recording their horizontal positions for stage movement and vertical positions for focusing. The brightfield images are taken at this point in time, certifying that the microscope will record sharp images at the set measurement points.

When the solutions have slowed down the syringes connected to the side inlets are replaced by ones filled with PBS buffer solution (at least 400 μl per syringe). The flow rate for these two syringes is increased to 14 μl/min for 15 minutes to make sure that the system is degassed, followed by another 10-15 minutes at 4 μl/min. The syringe connected to the middle inlet is replaced by one filled with 300 μl of the analyte, making sure that the syringe head is completely full of solution at the time of screwing in onto the syringe, since it is no longer possible to operate this particular pump at higher flow rates for degassing due to the limited amount of the analyte.

External light sources are then minimized around the microscope, and all three syringe pumps are operated at 4 μl/min until the analyte reaches the intersection of the device. This is monitored at an exposition rate of 300 ms and with 4x gain. When the analyte has reached the intersection its flow rate is reduced to 1 μl/min. The fluorescent images are recorded 10-15 minutes later at an exposition rate of 1-5 s (depending on signal strength) and with 1x gain. The temperature around the stage is recorded immediately after the fluorescent images.

The fluorescent and brightfield images are recorded at the same positions, which means that they can be easily overlapped in the microscope’s software. Since it is not feasible to attach the device to the stage in such a manner that the main channel is perfectly horizontal on the recorded images, they must be rotated by ± 2°. The intensity profiles are recorded along vertical vectors placed over the images, specifically where the signal-to-noise ratio is maximal. Fluorescent and brightfield profiles are recorded in pairs along the same vector. The distance of these vectors from the markers of the measurement points is also recorded. Finally, one pair of profiles is exported for each measurement point into distinct tabs of an Excel file, the first tab of which contains every other piece of recorded information that is relevant for the analytic software.

### The analytic software package

The measured experimental data is processed with a software package consisting of custom scripts and functions written in MATLAB. The main script takes a single Excel file containing experimental data as input. It omits any specified measurement points from the evaluation, e.g. the last profile should be omitted due to the laminar flow being disturbed by some sort of contaminant stuck between the last two measurement points. The following information are imported from the input file: The width and height of the main channel (μm), the thickness of its side walls as displayed in the brightfield recording (μm), the flow rate of the analyte and the buffer solutions (μl/min), the temperature around the stage (K), the viscosity of the analyte (Pa*s), the exposition time (s), the position of each profile, relative to the origin point at the intersection (μm), the fluorescent and brightfield intensity profiles (1). The script calculates the average velocity of particles:

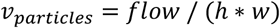

The average time it took for particles to reach each measurement point after reaching the origin point at the intersection is calculated by dividing the distance of the profile from the origin point by the average velocity above. After that, the average time particles spent between consecutive measurement points are yielded by simple subtractions. Since all particles can be assumed to have a hydrodynamic radius between 1 nm and 2 μm, it is possible to calculate the minimum and maximum incline for the linear function fitted to the time-variance function (see Basic principles of the experimental method).

In case the recorded images had to be rotated to align them horizontally the intensity profiles fall to zero at the edges where the vectors along which they were taken include points outside the images. A custom cropping function is called to remove these outside sections. They are recognized by defining a band around the brightfield profile as shown below. The cropping function starts from the two ends of the brightfield profiles and removes each point until it comes across one the intensity of which is higher than the lower limit of the band.

After cropping a custom normalization function is called that brings the signals measured outside the microfluidic channel to the same baseline, and then it aligns that baseline to the (intensity = 0) axis as well as centering the profiles around the fluorescent intensity maximum. All these steps require the positions of the channel’s sidewalls, in order to distinguish data points inside it from one outside it. This process starts with finding the position of the fluorescent maximum, from which the function iterates through data points in both directions in the associated brightfield profile until it comes across either the intensity minimum or maximum in that portion of the profile, whichever is further away from the mean value of the brightfield intensities. Since the cropping function previously removed data points outside of the recorded images, the only two sections in each brightfield profile where the intensity significantly deviates from the mean value are the two side walls. Based on the wall thickness defined on the input file, the function excludes sections around these two points, dividing each profile into three sections: the section inside the channel, the right-hand side external section and the left-hand side external section, defined relative to the flow direction. Each brightfield profile is multiplied with a coefficient to align it with the 200-intensity line, which is the most common baseline based on observations. The coefficient is simply the ratio of 200 and the mean value of data points inside the channel. In the case of fluorescent profiles normalization means a two-step process. The first step is aligning the baseline of each profile to the standard 200, same as before. The goal of this step is to remove the differences between profiles resulting from the slight changes in external noise. The second step is to multiply each profile with a coefficient so that all of them have the same area under their curves. The coefficient in this case is the ratio of the largest integral value and the integral of the current profile. These integrals ignore the sections outside the channel. Finally, the baseline of each fluorescent profile is aligned with the (intensity = 0) axis by subtracting the mean value of outside data points from the entire profile and is centered on the position of its peak value, a horizontal shift applied to the associated brightfield profile as well. The position of the peak value is often different from what would be the center of the curve; however, this slight discrepancy is nullified in the next step.

Each profile is then fitted with a Gaussian function or the linear combination of up to eight Gaussian functions, using MATLAB’s “prepareCurveData” and “fit” functions. The parameters of the fitted functions are limited as follows: the amplitude of the curve must not be negative, the center of the curve must be between -5 μm and 5 μm (the peak value is always within 5 μm of the center of the curve based on observations), and the standard deviation of the curve must be positive but not larger than 100, an upper limit selected because it has been observed to yield the best approximations for samples with known particle sizes. The limits of the standard deviation are adjusted for each profile based on the maximal and minimal changes theoretically possible between measurement points, based on the maximum and minimum incline calculated at the beginning of this section. This ensures that the fitted curves widen as we progress from the first measurement point towards the last, which is not guaranteed otherwise due to lingering artefacts. All fits are initiated from multiple randomized positions until a better fit could not be found for five consecutive attempts, evaluated based on RMSE. Increasing this number over five did not yield significant improvements in the RMSE of the fitted curves.

The custom function that handles the fitting of Gaussian curves, including the “prepareCurveData” and “fit” functions, generates 3D tensors from which the Gaussian coefficients (amplitude, center, standard deviation) and the “Goodness of Fit” metrics (RMSE, SSE, R-square, Adjusted R-square) are extracted for each selected order of fit from one to eight, and for each measurement point. For example, if the experimental data consists of ten fluorescent profiles, and each of those profiles were fitted with the linear combination of eight Gaussian functions, there will be eight sets consisting of ten Gaussian functions each, where each function belongs to a different measurement point, and their standard deviations, and thus their variances, form a linear, monotonically increasing function in time. Each of these monotonically increasing functions represent a particle size that is supposedly present within the measured analytes solution, and the diffusion coefficient associated with this particle size is one fourth of the incline of the associated monotonically increasing function. So, the final step the analytic software carries out before displaying the results is to fit a linear function to the monotonic function that displays the variances of the Gaussian function within the same set against the estimated average time particles spent between measurement points (see Fig. 1.). Then, the diffusion coefficients can be determined, and particle radii can be approximated (see Basic principles of the experimental method).

## Author Contributions Statement

Z.G. and A.L.S. designed the study; M.L., A.J.L., and A.L.S. developed the device; E.N.K., S.V., and E.A.J. prepared the samples; A.L.S., E.A.J., and C.I.P. carried out the measurements; A.L.S. developed the analytic software; Z.G., A.L.S., and E.N.K. analyzed the data; all authors contributed to data interpretation, drafting the article and approved the final version.

## Acknowledgements

This work was supported by a grant from the National Research, Development and Innovation Office under grants TKP2021-EGA-42 and OTKA 137947.

The authors thank Tünde Juhász from the Biomolecular Self-assembly Research Group at the HUN-REN Research Centre for Natural Sciences for her kind help with the DLS measurements. We also thank Márton Iványi for his involvement in the production of FITC-labelled Drebrin samples. The kind help of Dr. Kristóf Iván for his suggestions on the manuscript is also acknowledged. This work was supported by a grant from the National Research, Development and Innovation Office under grants TKP2021-EGA-42 and OTKA 137947.

## Notes

### Competing Interest Statement

The authors have declared no competing interest.

### Summary of Updates

Additional experiments have been included and the purpose of our work has been clarified in the introduction section.

